# Mouse-embryo model derived exclusively from embryonic stem cells undergo neurulation and heart development

**DOI:** 10.1101/2022.08.01.502371

**Authors:** Kasey Y.C. Lau, Hernan Rubinstein, Carlos W. Gantner, Gianluca Amadei, Yonatan Stelzer, Magdalena Zernicka-Goetz

**Affiliations:** Department of Physiology, Development and Neuroscience, University of Cambridge; Cambridge CB2 3EG, UK; Department of Molecular Cell Biology, Weizmann Institute of Science; 7610001 Rehovot, Israel; California Institute of Technology, Division of Biology and Biological Engineering; 1200 E. California Boulevard, Pasadena, CA 91125, USA

## Abstract

Several *in vitro* models have been developed to recapitulate mouse embryogenesis solely from embryonic stem cells (ESCs). Despite mimicking many aspects of early development, they fail to capture the interactions between embryonic and extraembryonic tissues. To overcome this difficulty, we have developed a mouse ESC-based *in vitro* model that reconstitutes the pluripotent ESC lineage and the two extra-embryonic lineages of the post-implantation embryo by transcription factor-mediated induction. This unified model recapitulates developmental events from embryonic day 5.5 to 8.5, including gastrulation, and formation of the anterior-posterior axis, brain, a beating heart structure, and the development of extraembryonic tissues, including yolk sac and chorion. Comparing single-cell RNA sequencing from individual structures with time-matched natural embryos identified remarkably similar transcriptional programs across lineages, but also showed when and where the model diverges from the natural program. Our findings demonstrate an extra-ordinary plasticity of ESCs to self-organize and generate a whole embryo-like structure.

## Introduction

At the time of implantation, the mouse blastocyst comprises three lineages: the epiblast (EPI), the trophectoderm (TE), and the primitive endoderm (PE) that will give rise to the embryo proper, the placenta, and the yolk sac respectively. By using stem cells derived from these lineages, several *in vitro* models have been developed to recapitulate various events of post-implantation development. One approach has been to take solely mouse embryonic stem cells (ESCs) and by applying exogenous stimuli induce them to establish anterior-posterior polarity (ten Berge *et al*., 2008) and mimic basic body axis formation, and aspects of gastrulation, somitogenesis, cardiogenesis and neurulation (Van Den Brink *et al*., 2014; Turner *et al*., 2017; Beccari *et al*., 2018; Rossi *et al*., 2020; Veenvliet *et al*., 2020; Xu *et al*., 2021). Such so-called “gastruloids” are a powerful system and demonstrate the ability of ESCs to be directed into complex developmental programs. However, these systems fail to capture the entire complexity of signalling and morphological events along the complete body axes. This is largely because they fail to recapitulate the spatio-temporal interplay of signalling pathways between embryonic and extraembryonic tissues, which is crucial to pattern the post-implantation mouse embryo. Consequently, they do not represent complete embryonic structures and lack the overall morphological resemblance to natural post-implantation mouse embryos.

We have therefore adopted a second approach to fully model the post-implantation mouse embryo by promoting assembly of mouse <u>E</u>SCs with either extraembryonic trophoblast stem cells (<u>T</u>SCs), to direct formation of a post-implantation egg cylinder showing appropriate posterior development (Harrison *et al*., 2017), or a mixture of TSCs and extraembryonic endoderm (<u>X</u>EN) stem cells to generate “ETX” embryos that develop anterior and posterior identity and initiate gastrulation movements (Sozen *et al*., 2019). Subsequently, by replacing XEN cells with ESCs harbouring inducible Gata4 expression (iGata4 ESCs), it proved possible to generate XEN cells at an earlier stage of development which could contribute to iETX embryos that were fully able to recapitulate gastrulation movements (Amadei *et al*., 2021). One remaining complication of the iETX embryo model is that TSCs and ESCs require different culture media, necessitating the use of undefined culture conditions and increasing the difficulty of developing embryoids in the laboratory. Therefore, we have developed an entirely ESC-based *in vitro* model that reconstitutes the three fundamental cell lineages of the natural post-implantation mouse embryo through transcription factor-mediated reprogramming. In addition to replacing XEN cells with induced XEN cells, we now further substitute TSCs with ESCs that transiently overexpress *Cdx2* upon doxycycline induction. We show that such induced TSCs could effectively replace TSCs to form embryo-like structures which we termed “EiTiX-embryoids”. These EiTiX-embryoids undergo development from pre-gastrulation stages to neurulation stages, developing headfolds, brain, a beating heart structure, and extraembryonic tissues, including a yolk sac and chorion. In agreement with the similar overall morphology, our single cell, single structure analysis reveals a robust recapitulation of cell states spanning both embryonic and extraembryonic lineages, with strikingly little variation in overall gene expression program in these states. Yet, our approach also demonstrates that *Cdx2*-expressing cells can contribute to the chorion but not the ectoplacental cone lineage in the extraembryonic ectoderm compartment.

## Results

### *Cdx2*-induced ESCs self-assemble with *Gata4*-induced ESCs and ESCs into post-implantation-like mouse embryoids

Cdx2 is a key transcription factor driving TE development and its overexpression leads ESCs to transdifferentiate into TSC-like cells (Niwa *et al*., 2005). To determine whether *Cdx2*-expressing ESCs could replace TSCs in generating ETiX-embryoids, we generated a transgenic ESC line carrying a doxycycline (Dox)-inducible *Cdx2* gene. The resulting clones of i*Cdx2*-ESCs showed a 100- to 200- fold increase in *Cdx2* mRNA expression after 6 hours of Dox induction (Figure S1A). From the four clones we tested, we selected the clone with the highest level of *Cdx2* overexpression for subsequent experiments. This clone showed a substantial upregulation of both *Cdx2* mRNA (Figure S1B) and protein, as detected by qRT-PCR and immunofluorescence respectively, after 6 hours of induction (Figure S1C). To assess the long-term effect of Cdx2 overexpression on cell fate, we compared three different types of cell aggregates: either induced i*Cdx2* ESCs, uninduced i*Cdx2* ESCs; or TSCs (Figure 1A). After three days, we observed a significant upregulation of the TSC marker, Eomes, and downregulation of the ESC marker, Oct4, in the aggregates of induced i*Cdx2* ESCs (Figures 1B, 1C, S1D and S1E). Transcripts of the TSC markers *Elf5*, *Eomes* and *Gata3* were also upregulated in the induced i*Cdx2* ESC aggregates (Figures S1F-H). Together, these findings suggest that upon *Cdx2* overexpression, i*Cdx2* ESCs lose their ESC identity and acquire TSC-like cell fate.

**Figure 1.**
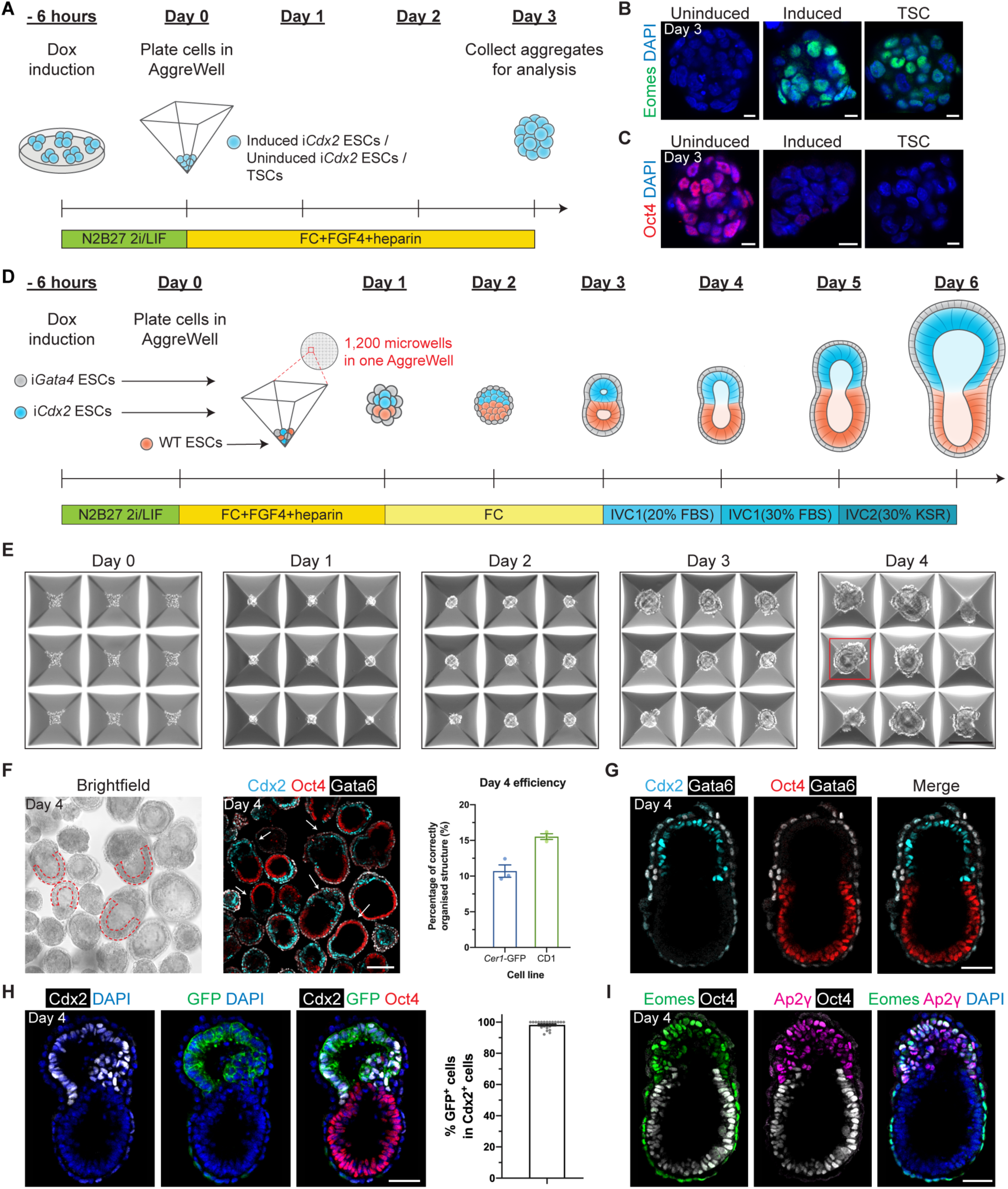
*Cdx2*-induced ESCs self-assemble with *Gata4*-induced ESCs and ESCs into post-implantation-like mouse embryoids. (**A**) Schematic of the formation of cell aggregates in AggreWells. Day 3 i*Cdx2* ESC aggregates show elevated Eomes expression (**B**) and downregulated Oct4 (**C**) upon induction of *Cdx2*. (**D**) Schematic of EiTiX-embryoid generation. (**E**) Representative brightfield images of structures developing in AggreWells from Day 0 to Day 4. A structure resembling the early post-implantation mouse embryo can be seen on Day 4 in the well outlined in red. (**F**) All structures in the combined microwells from one AggreWell were collected at Day 4 and stained to reveal Cdx2 (cyan), Oct4 (red) and Gata6 (white). Arrows indicate structures considered to exhibit correct organisation. Such cylindrical structures with two cellular compartments and an epithelialized EPI-like cell layer (red dashed outline) were selected under brightfield microscopy. The efficiency of obtaining organised structures using *Cer1*-GFP lines (combined *Cer1*-GFP unmodified; *Cer1*-GFP *Gata4* inducible ESC; and CAG-GFP *Cdx2* inducible ESC lines) or CD1 lines (combined CD1 unmodified; CD1 *Gata4* inducible ESC; and CAG-GFP *Cdx2* inducible ESC lines) is shown. (**G**) Day 4 EiTiX-embryoids stained to reveal Cdx2 (cyan), Oct4 (red) and Gata6 (white). (**H**) Day 4 EiTiX-embryoid stained to reveal Cdx2 (white), GFP (green) and Oct4 (red). The percentage of the Cdx2-positive cells that are also GFP-positive is shown. n = 49 structures. (**I**) Day 4 EiTiX-embryoids stained to reveal Eomes (green), Ap2γ (magenta) and Oct4 (white). n = 35/35 structures are positive for both Eomes and Ap2γ. All experiments were performed minimum 3 times. Scale bars: 10µm, (B-C); 150µm, (E-F); 50µm, (G-I).

We then asked whether induced i*Cdx2* ESCs could replace TSCs in generating embryoids when aggregated with wild-type (WT) ESCs and *Gata4* induced ESCs. To this end, we adapted our previously described protocol (Amadei *et al*., 2021) by inducing expression of *Cdx2* and *Gata4* by treating both i*Cdx2*- and i*Gata4*-ESC lines for 6 hours with doxycycline before combining them with WT ESCs in AggreWell plates (Figure 1D). Over the course of four days, we observed drastic morphological changes of the resulting cell aggregates such that by Day 4 we could observe structures that resembled post-implantation embryos, which naturally comprise EPI and TE-derived extra-embryonic ectoderm (ExE) compartments surrounded by visceral endoderm (VE) (Figure 1E). The random nature of the interactions of the three cell types results in a variety of structures (Figure S1I) but we optimized the efficiency of correct structure formation by adding FGF4 and heparin during the first 24 hours after plating and by doubling the number of i*Cdx2* ESCs seeded from 16 to 32 per microwell (Figures 1D and 1F). When using the *Cer1*-GFP ESC line (which monitors formation of the anterior signalling center, anterior VE (AVE)) and the CD1 ESC line, the efficiency of correct structure formation was 10.7% and 15.5%, respectively (Figure 1F). For all subsequent experiments, we selected structures with morphology resembling natural mouse embryos on Day 4 (Figures 1F and 1G, see Methods for inclusion criteria). Expression of the constitutive membrane GFP marker in i*Cdx2* ESCs showed that they had given rise to the Cdx2-positive cells that correctly localised within the ExE-like compartment (Figure 1H). We also detected the expression of other ExE markers including Ap2γ and Eomes in the putative ExE compartment, suggesting downstream TSC markers were also upregulated after *Cdx2* overexpression (Figure 1I). Finally, we compared the dimensions of EiTiX-embryoids, E5.5 mouse embryos and Day 4 iETX-embryoids and found that EiTiX-embryoids were most similar to E5.5 embryos (Figures S1J-M). Together, our findings show that i*Cdx2* ESCs can replace TSCs to generate post-implantation embryo-like structures expressing canonical lineage markers. Since <u>T</u>SCs were replaced by i*Cdx2* ESC, we termed the structures “EiTiX-embryoids.”

### EiTiX-embryoids establish an anterior-posterior axis and undergo gastrulation

We then asked whether EiTiX-embryoids could recapitulate key events of post-implantation development. We first asked whether the critical anterior signalling center can be formed, which breaks mouse embryo symmetry and establishes the anterior-posterior axis (Thomas and Beddington, 1996). This center first appears as the distal visceral endoderm (DVE) at the distal tip of the egg cylinder before migrating to the anterior side of the egg cylinder to become the anterior visceral endoderm (AVE), which is characterised by the expression of Cer1, Lefty1 and Dkk1 (Figure 2A). To this end, we formed EiTiX-embryoids using i*Gata4* ESCs with a *Cer1*-GFP reporter and observed the co-expression of GFP with Dkk1 or Lefty1 in EiTiX-embryoids at Day 4 and Day 5 (Figures 2B, S2A and S2B). To follow the development of the *Cer1*-GFP- positive domain, we determined the extent of AVE anterior migration (see Methods) and binned the measurements into three groups: ‘proximal’, >67% migration; ‘lateral’, 33-67% migration, and ‘distal’, <33% migration. Anterior migration of the AVE was evident from the higher proportion of Day 5 EiTiX-embryoids with proximal *Cer1*-GFP and Dkk1 expression than in Day 4 EiTiX-embryoids, while the distribution of Lefty1-positive domain remained similar (Figures 2C and S2C).

**Figure 2.**
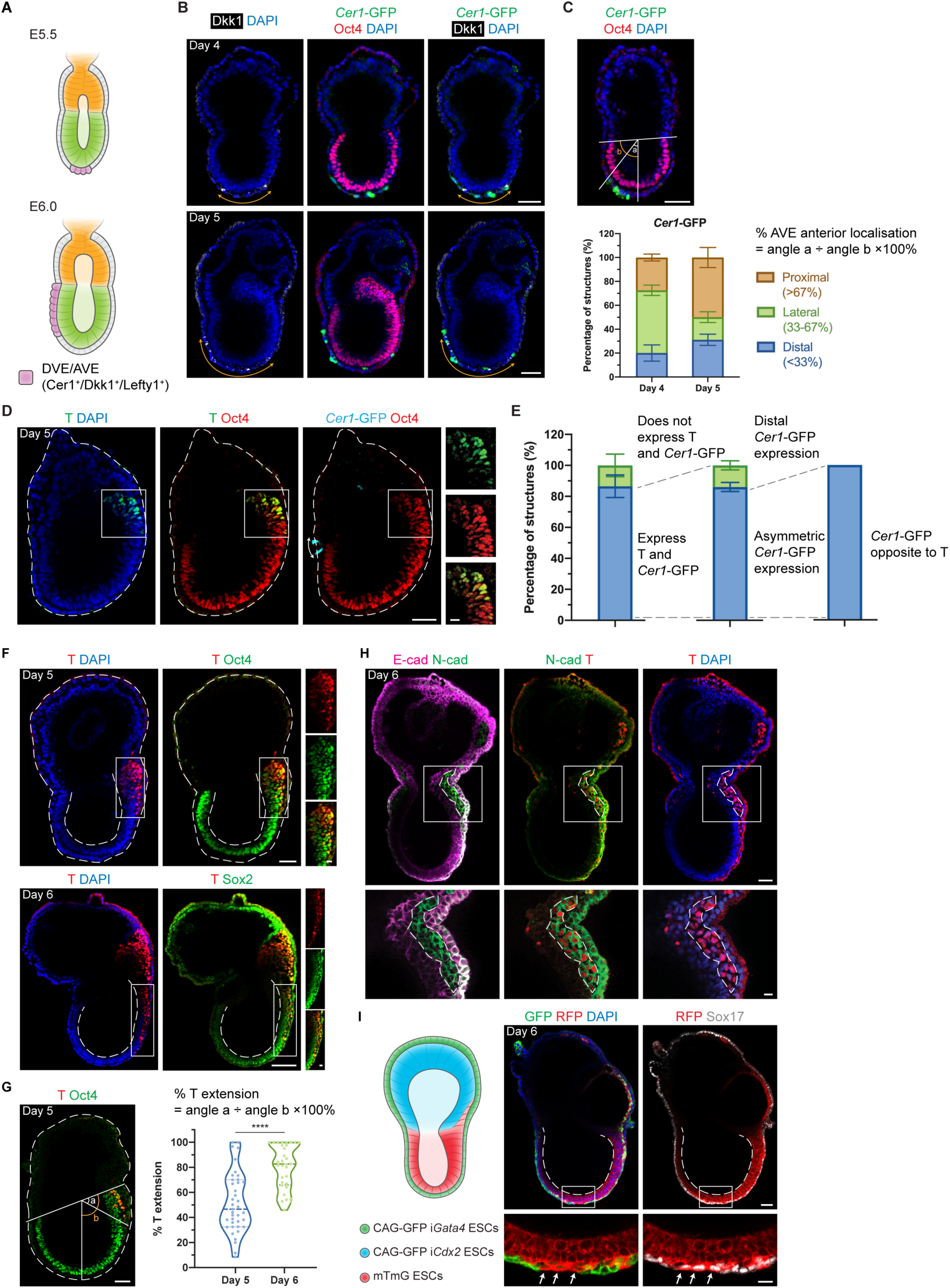
EiTiX-embryoids establish an anterior-posterior axis and undergo gastrulation. (**A**) Schematic showing the position of DVE/AVE in E5.5 and E6.5 mouse embryos. (**B**) Day 4 and Day 5 EiTiX-embryoids stained to reveal *Cer1*-GFP (green), Oct4 (red) and Dkk1 (white). Orange double-headed arrows indicate Dkk1-positive domains. (**C**) Localisation of *Cer1*-GFP in Day 4 and Day 5 EiTiX-embryoids (See Materials and Methods for quantification method). n = 57 Day 4 structures; 45 Day 5 structures. (**D**) Day 5 EiTiX-embryoids stained to reveal T (green), Oct4 (red) and *Cer1*-GFP (cyan). Double-headed arrow, *Cer1*-GFP-expressing domain; white box outline, T- and Oct4- positive domain. (**E**) Percentages of Day 5 EiTiX-embryoids showing 1) expression of both *Cer1*-GFP and T in same structure, 2) asymmetric *Cer1*-GFP expression, and 3) T expression on the opposite side from *Cer1*-GFP. n = 42 structures. (**F**) Day 5 and Day 6 EiTiX-embryoids stained to reveal T (red) and Oct4 (green) or Sox2 (green). White boxes enclose T-positive domain while dotted lines outline the structure and the lumen of ES compartment. n = 39/42 Day 5 structures and 32/32 Day 6 structures. (**G**) Percentage of T extension in Day 5 and Day 6 EiTiX-embryoids. See Materials and Methods for quantification method. n = 39 Day 5 structures and 32 Day 6 structures. ****p < 0.0001. (**H**) Day 6 EiTiX-embryoid stained to reveal E-cadherin (magenta), N-cadherin (green) and T (red). Dotted line indicates T- and N-cadherin-positive domain. n = 14/21 structures with N-cadherin upregulation and E-cadherin downregulation from 4 experiments. (**I**) Schematic of EiTiX-embryoids using CAG-GFP i*Gata4* ESCs (with membrane GFP) and mTmG ESCs (with membrane tdTomato) to construct the VE-like layer and EPI-like compartment, respectively. Day 6 EiTiX-embryoid stained to reveal GFP (green), RFP (red) and Sox17 (white). Dotted line indicates the lumen of ES compartment while arrows mark definitive endoderm-like cells intercalated into the VE-like layer. n = 8/12 structures. All experiments were performed minimum 3 times. Scale bars: 50µm; 15µm (zoomed).

In the natural post-implantation embryo, the AVE is critical to restrict primitive streak formation to the posterior EPI through the secretion of Nodal and Wnt inhibitors (Stower and Srinivas, 2017). We therefore asked whether these events could be recapitulated in EiTiX-embryoids and analyzed the expression of Brachyury (T), a primitive streak marker, in relation to the *Cer1*-GFP domain at Day 5. We found that 86.7% of Day 5 EiTiX-embryoids expressed *Cer1*-GFP and T, and of these, 86% showed opposed *Cer1*-GFP and T expression (Figures 2D and 2E). Similarly, we found that 94.7% of structures with asymmetric AVE expression of Cer1-, Dkk1-, or Lefty1 showed expression of the primitive streak marker, Eomes, on the opposite side (Figures S2D-F). Thus, EiTiX-embryoids correctly establish both the AVE and primitive streak, recapitulating anterior-posterior patterning as in natural post-implantation embryos.

After establishment of the anterior-posterior axis and onset of gastrulation in the posterior EPI, the primitive streak extends to the distal end of the egg cylinder (Bardot and Hadjantonakis, 2020). Accordingly, as EiTiX-embryoids developed, we could detect T- and Oct4-positive cells at the posterior end of the EPI-like compartment on Day 5, that had extended to the distal-most part of the egg cylinder on Day 6 (Figure 2F). To quantify the percentile extension of this T-positive domain, we measured the angle between the posterior boundary of the Oct4-positive domain and the most anterior T-positive cells (Figure 2G, angle a) divided by the angle subtended by the Oct4-positive domain boundary and the distal tip (angle b), where 100% indicates complete extension. This showed the degree of extension approached its fullest extent at Day 6 (Figure 2G). As cells egress from the EPI to form the primitive streak, they undergo epithelial-to-mesenchymal transition, downregulating E-cadherin and upregulating N-cadherin (Arnold and Robertson, 2009). We observed a T-positive domain that had robust N-cadherin expression but unlike surrounding cells, did not express E-cadherin in 66.7% of EiTiX-embryoids at Day 6 (Figure 2H).

As development progresses, the primitive streak undergoes further specification to produce a range of cell types, including axial mesendoderm and definitive endoderm (Bardot and Hadjantonakis, 2020; Scheibner *et al*., 2021). In Day 6 EiTiX-embryoids, we could detect the presence of T- and Foxa2-positive cells identifying axial mesendoderm (93.8%), as well as Foxa2- and Sox17-positive cells identifying definitive endoderm (88.2%) (Figures S2G and S2H). In the natural mouse embryo, the EPI-derived definitive endoderm gradually displaces and intercalates with the VE which covers the egg cylinder (Kwon, Viotti and Hadjantonakis, 2008). To visualise whether such endoderm intercalation takes place in EiTiX-embryoids, we used ESCs expressing membrane tdTomato (mTmG ESCs) to generate the EPI-like compartment and i*Gata4* ESCs expressing membrane GFP (CAG-GFP i*Gata4* ESCs) to generate the VE-like layer in the embryoids. We observed a discontinuous GFP-positive cell layer interspersed with RFP-positive cells (Figure 2I, 66.7%). These cells also expressed Sox17 which is a critical factor for endoderm specification and for the egression of definitive endoderm cells into the VE (Viotti, Nowotschin and Hadjantonakis, 2014). Thus, the intercalation of VE into the definitive endoderm is recapitulated in the EiTiX-embryoids.

### Day 6 EiTiX-embryoids capture major cell types of gastrulation

After finding that Day 6 EiTiX-embryoids could capture numerous processes of gastrulation, we sought to understand the overall cell type composition of these gastrulating EiTiX-embryoids in comparison to natural embryos. We utilized a recently established time-resolved model of mouse gastrulation consisting of ∼68,000 single cells derived from 287 individually processed embryos spanning egg cylinder stage to early somitogenesis (Figure 3A, Mittnenzweig *et al*., 2021), which: (i) enables a quantitative evaluation of transcriptional states; and (ii) describes the natural flux of embryonic and extraembryonic lineage differentiation, thus allowing analysis of cell state composition within individual structures. We generated Day 6 EiTiX-embryoids by combining i*Cdx2* ESCs with constitutive membrane GFP-expression (CAG-GFP), unlabelled WT ESCs and i*Gata4* ESCs carrying the *Cer1*-GFP reporter. GFP signals confirmed the appearance of the ExE-like compartment and the AVE-like domain in EiTiX-embryoids (Figure 3B). Next, we performed single-cell RNA sequencing (scRNA-seq) on 14 individual EiTiX-embryoids using MARS-seq by index-sorting into barcoded 384-well plate as previously reported (Mittnenzweig *et al*., 2021).

**Figure 3.**
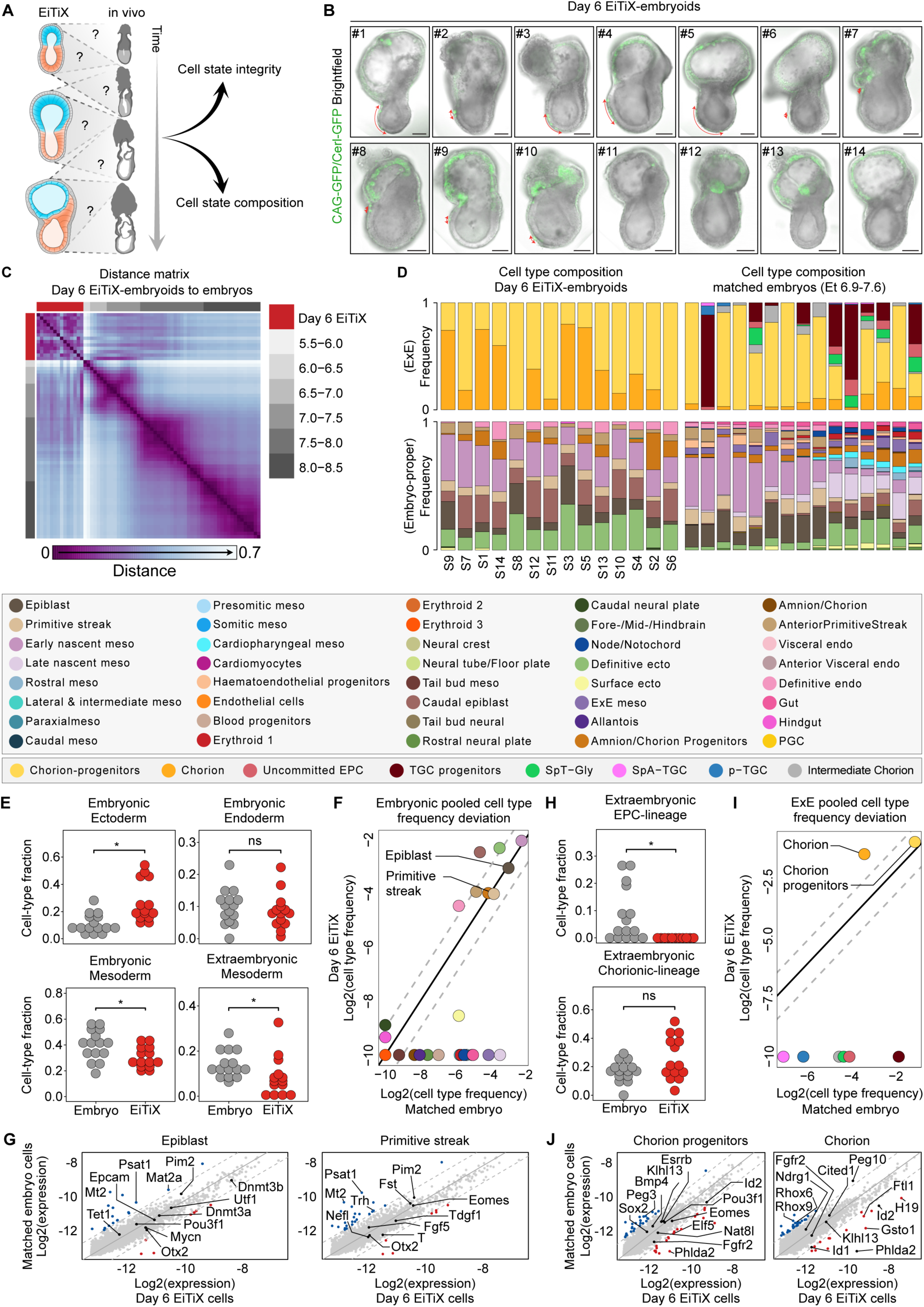
Day 6 EiTiX-embryoids capture major cell types of gastrulation. (**A**) Evaluation of cell state integrity and cell type composition of Day 6 EiTiX-embryoids using our recently established time-resolved model of mouse gastrulation. (**B**) Brightfield images of Day 6 EiTiX-embryoids (n = 14, annotated as S#1-14) collected for single-structure, single-cell RNA sequencing, overlaid with GFP expression (*Cer1*-GFP reporter in VE-like layer and membrane CAG-GFP from i*Cdx2* ESCs in ExE-like compartment). Red arrow indicates *Cer1*-GFP expression. Scale bar: 100µm. (**C**) Embryo-embryo cell type composition similarity matrix. Natural embryos are annotated based on embryonic age groups (grey), Day 6 EiTiX-embryoids (red). (**D**) Cell type composition bars of individual Day 6 EiTiX-embryoids (left) and matched natural embryos (right, annotated according to the key shown below). (**E, H**) Comparison of major embryonic germ layers frequency (E) and ExE lineages frequency (H) between Day 6 EiTiX-embryoids and matching natural embryos. Medians of frequencies were compared using a Wilcoxon-Mann-Whitney rank sum test after down sampling of cell state specific cells to corresponding number of Day 6 EiTiX cell state specific cells (i.e. for each cell state individually). q Values were calculated from p values according to the Benjamini-Hochberg procedure. ns, not significant; *, q value < 0.05. Major germ layers - *Embryonic ectoderm*; Forebrain/Midbrain/Hindbrain, Rostral neural plate, Surface ectoderm, Caudal neural plate, Definitive ectoderm. *Embryonic endoderm*; Definitive endoderm, Gut, Hindgut, Visceral and Anterior Visceral endoderm. *Embryonic mesoderm*; Tail bud-, Early and Late nascent-, Caudal-, Presomitic-, Somitic-, Paraxial-, Rostral-, Cardioparyngeal- and Lateral & intermediate-mesoderm. *ExE Mesoderm*; Amnion/Chorion progenitor, Amnion/Chorion, Allantois and ExE mesoderm. *EPC-lineage*; SpT-Gly, TGC progenitors, uncommitted EPC, pTGC and SpA-TGCs. *Chorion-lineage*; intermediate ExE, Chorion progenitors and Chorion. (**F, I**) Pooled embryonic (F) and ExE (I) cell type frequencies comparison between Day 6 EiTiX-embryoids and matched natural embryos. (**G, J**) Bulk differential gene expression per cell type of Day 6 EiTiX cells against matched embryo cells in embryonic cell types (G, EPI and primitive streak) and ExE cell types (J, chorion progenitors and chorion). Dots represent individual genes. Color annotated dots mark genes with a two-fold change in expression (blue – above two-fold decrease in Day 6 EiTiX cells, red – above two-fold increase in Day 6 EiTiX cells).

A strategy for ranking embryos by K-nn similarities among their single-cell profiles identified high similarity between individual Day 6 EiTiX-embryoids, which cluster separately from natural embryos. In agreement with the morphological assessment of these embryoids, their overall transcriptional ranking was found to be most similar to E6.5-7.5 gastrulation stages (Figures 3C and S3A). Next, we constructed and annotated a transcriptional manifold of EiTiX-embryoids (see Methods). Remarkably, we found robust mapping to unmodified embryonic and extraembryonic cell states and did not detect any non-coherent transcriptional programs in the metacells of Day 6 EiTiX-embryoids, suggesting they conserve the transcriptional programs of the corresponding cell states in natural embryos (Figure S3B). Focusing first on embryonic cell state compositions, we observed a high degree of similarity among the 14 Day 6 EiTiX-embryoids, despite them having variable morphologies (Figure 3D). However, when compared to natural embryos from corresponding time bins (Methods), we found deviations from the natural program. First, we noted some of the lineages were not in synchrony. Specifically, while differentiation of nascent mesoderm to extraembryonic mesoderm, blood, and hematoendothelial progenitors resembled younger natural embryos (∼E6.5), both definitive ectoderm, tail-bud EPI, and amnion were overrepresented, resembling the composition of more advanced natural embryos (∼E7.5) (Figures 3D-F). Yet, highly similar gene expression profiles were indicated by the presence of cell states in comparable frequencies to those in natural embryos (Figures 3G and S3C-D).

Ectoplacental cone cells (EPC) were absent from Day 6 EiTiX-embryoids compared to respective natural embryos. In contrast, the chorion lineage, comprising both chorion progenitors and their differentiated progenies, was largely intact (Figures 3D, 3H, and 3I). Gene expression analysis showed both programs to be overall highly similar to the natural embryos (Figure 3J). It also revealed down-regulation of bona-fide chorion genes *Rhox6* and *Rhox*9, together with upregulation of *Id2*, consistent with a lack of proximal signals emanating from the EPC compartment. Taken together, our analysis identified remarkably similar transcriptional states between Day 6 EiTiX-embryoids and their natural counterparts, but it also revealed pausing in mesoderm differentiation and over-accumulation of posterior cell types, most likely reflecting alterations in synchronicity between lineages. We therefore next asked whether further culture of embryoids would enhance the synchronicity of lineage development.

### EiTiX-embryoids develop to late headfold stages with heart and chorion development

To assess the full developmental potential of EiTiX-embryoids, we next transferred them to a recently reported *ex utero* culture medium (EUCM) (Aguilera-Castrejon *et al*., 2021) (Figure 4A). We found that EiTiX-embryoids developed regions resembling headfolds, a beating heart, allantois and chorion over the next three days in culture and that they shared highly similar morphologies with E6.5 natural embryos cultured in EUCM (Figures 4B, movie S1 and S2). EiTiX-embryoids also developed a yolk sac-like membrane which enclosed the embryonic structures (Figure 4C). The efficiencies of EiTiX-embryoids progressing from Day 5 to 6, Day 6 to 7 and Day 7 to 8 were between 65.4% and 75% (Figure 4D). Successfully developed Day 8 EiTiX-embryoids have well defined structures resembling headfolds, heart and tailbud although we frequently observed enlarged heart structures (Figure S4A). The most commonly observed phenotypes of underdeveloped Day 8 EiTiX-embryoids include stunted overall development and impaired axial elongation to generate posterior structures (Figure S4B).

**Figure 4.**
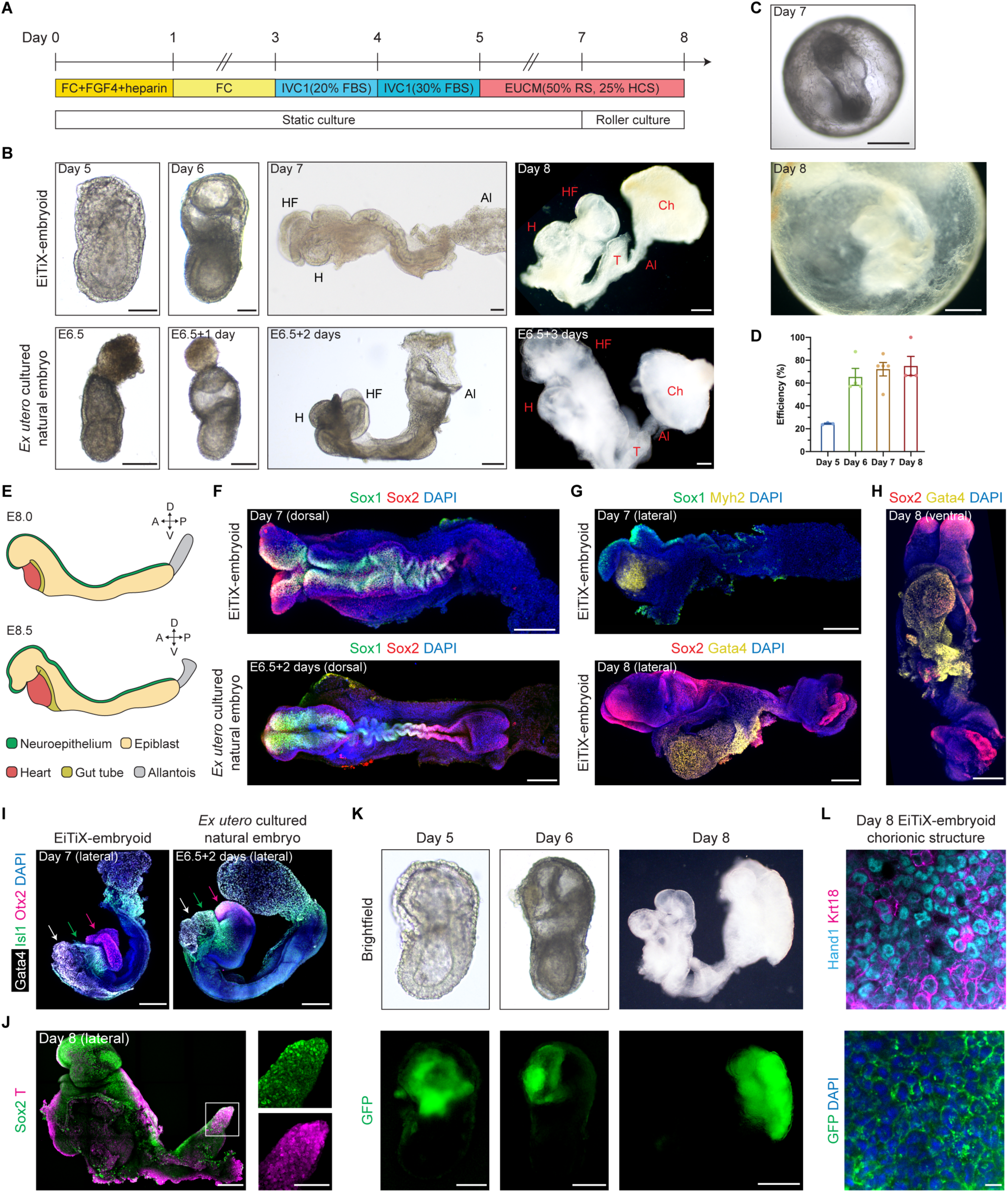
EiTiX-embryoids develop to late headfold stages with heart and chorion development. (**A**) Schematic showing culture conditions of EiTiX-embryoids to Day 8. (**B**) Representative brightfield images EiTiX-embryoids cultured from Day 5 to 8 and E6.5 natural embryo cultured *ex utero* for 3 days. Al: allantois, Ch: chorion, H: heart, HF: headfolds, T: tailbud. Scale bar, 100µm. (**C**) Brightfield images of Day 7 and Day 8 EiTiX-embryoids before dissecting yolk sac-like membrane. Scale bar, 200µm (Day 7); 500µm (Day 8). (**D**) Efficiency of EiTiX-embryoid progression from Day 4 to 5, Day 5 to 6, Day 6 to 7, and Day 7 to 8. (**E**) Schematic showing major cell types in E8.0 and E8.5 embryos. (**F**) Dorsal view of Day 7 EiTiX-embryoid and E6.5 natural embryo cultured *ex utero* for 2 days and stained to reveal Sox1 (green) and Sox2 (red). Scale bar, 200µm. (**G**) Lateral view of Day 7 and Day 8 EiTiX-embryoids stained to reveal neuroepithelial markers Sox1 (green) or Sox2 (red) and heart markers Myh2 or Gata4 (yellow). Scale bar, 200µm. (**H**) Ventral view of Day 8 EiTiX-embryoid stained to reveal Sox2 (red) and Gata4 (yellow), resembling the linear heart tube stage. Scale bar, 200µm. (**I**) Lateral view of Day 7 EiTiX-embryoid and E6.5 natural embryo cultured *ex utero* for two days stained to reveal heart marker Gata4 (white), pharyngeal mesoderm marker Isl1 (green), and forebrain marker Otx2 (magenta). Scale bar, 200µm. (**J**) Lateral view of Day 8 EiTiX-embryoids stained to reveal Sox2 (green) and T (red). Magnified panel showing co-expression in tailbud region (white square). Scale bar, 200µm; 100µm (zoomed). (**K**) Representative brightfield and GFP fluorescence image of Day 5 to 8 EiTiX-embryoids to track the contribution of CAG-GFP i*Cdx2* ESCs. Structures show GFP expression in chorion-like region. Scale bar, 50µm (Day 5); 200µm (Day 6); 500µm (Day 8). (**L**) Dissected chorionic structure from Day 8 EiTiX-embryoid stained to reveal GFP (green), Hand1 (cyan) and Keratin18 (magenta). Scale bar, 100µm.

Similar to natural E8.0 embryos and E6.5 embryos cultured *ex utero* for 2 days, the neuroepithelium markers Sox1 and Sox2 were expressed along the anterior-posterior axis of Day 7 EiTiX-embryoids (Figures 4E and 4F), indicative of neurulation. Interestingly we observed twisting of the neural tube-like region in both Day 7 EiTiX-embryoid and *ex utero* cultured embryo, suggesting that this could be a defect of *ex utero* culture. The heart markers Myh2 and Gata4 were expressed below the headfolds (Figure 4G), and a ventral view of the Gata4-expressing heart region revealed a morphology that resembled the linear heart tube (Figure 4H). Importantly, the anterior region of the headfolds showed robust Otx2 expression, indicating development of the forebrain (Figures 4I and S4C). We also detected the expression of Islet1 (Isl1), a pharyngeal mesoderm marker, between the Gata4-expressing heart region and Otx2-expressing forebrain region, recapitulating the expression pattern in the *ex utero* cultured embryo (Figures 4I and S4C). At the posterior end of the body axis, we observed robust co-expression of Sox2 and T at the region resembling the tail bud, which identifies the neuromesodermal progenitor population (Figure 4J).

As the neurulating EiTiX-embryoids arise from ESC and two different types of induced extraembryonic ESC types, we wished to determine the extent of extraembryonic tissue development. We detected chorion, an ExE-derived tissue that forms part of the placenta, and chorion progenitors in Day 6 EiTiX-embryoids by scRNA-seq (Figure 3D) and could observe a region that resembled the chorion in Day 8 EiTiX-embryoids (Figure 4K). To ask whether this region was derived from i*Cdx2* ESCs, we generated Day 8 EiTiX embryos using i*Cdx2* ESCs constitutively expressing membrane associated GFP and combined these with unlabelled WT and i*Gata4* ESCs. We observed membrane-associated GFP in the ExE region in Day 5 and Day 6 EiTiX-embryoids and at Day 8, the membrane GFP was exclusively found in the region resembling the chorion (Figure 4K). Further examination of this latter region showed co-expression of the chorion markers, Hand1 and Keratin18 (Figure 4L).

Taken together, these results indicate that EiTiX-embryoids have the remarkable ability to develop to headfold stages. They not only give rise to advanced embryonic structures such as neuroepithelium, a beating heart and mesodermal populations, but importantly, also develop extraembryonic tissues including yolk sac and chorion. The tracking of membrane GFP-positive i*Cdx2* ESCs further confirmed i*Cdx2* ESCs can effectively develop into chorion, an ExE-derived tissue, demonstrating that we could generate an embryoid with embryonic and extraembryonic tissues entirely from ESCs.

### Cell state and composition analysis of neurulating embryoids using scRNA-seq

To undertake a comprehensive analysis of cell state integrity and composition in neurulating embryoids, we collected four Day 8 EiTiX-embryoids for scRNA-seq (Figure 5A). Transcriptional similarity analysis showed Day 8 EiTiX-embryoids cluster separately from natural embryos, but overall most resemble E8.0-8.5 stages (Figure 5B). Analysis of cell state composition confirmed the high similarity between individual embryoids. In addition, Day 8 embryoids displayed high synchronicity between lineages and exhibited advanced cell states consistent with late head fold stages (Figures 5C and S5A). Quantitative analysis of cell-state frequency deviations identified depletion of tail-bud cell types and hematoendothelial progenitors. Furthermore, consistent with the enlarged heart structure (Figure S4A), we found an over-representation of cardiomyocytes (Figure 5D). We note that these deviations must be viewed cautiously, given the low frequency associated with some of these cell types. For example, although presomitic mesoderm was not present in Day 8 EiTiX-embryoids, a bona-fide somitic mesoderm population could be detected, suggesting that this progenitor population was merely not sampled. Finally, we detected similar gene expression patterns in high-frequency cell states compared to controls (Figures 5E and S5B).

**Figure 5.**
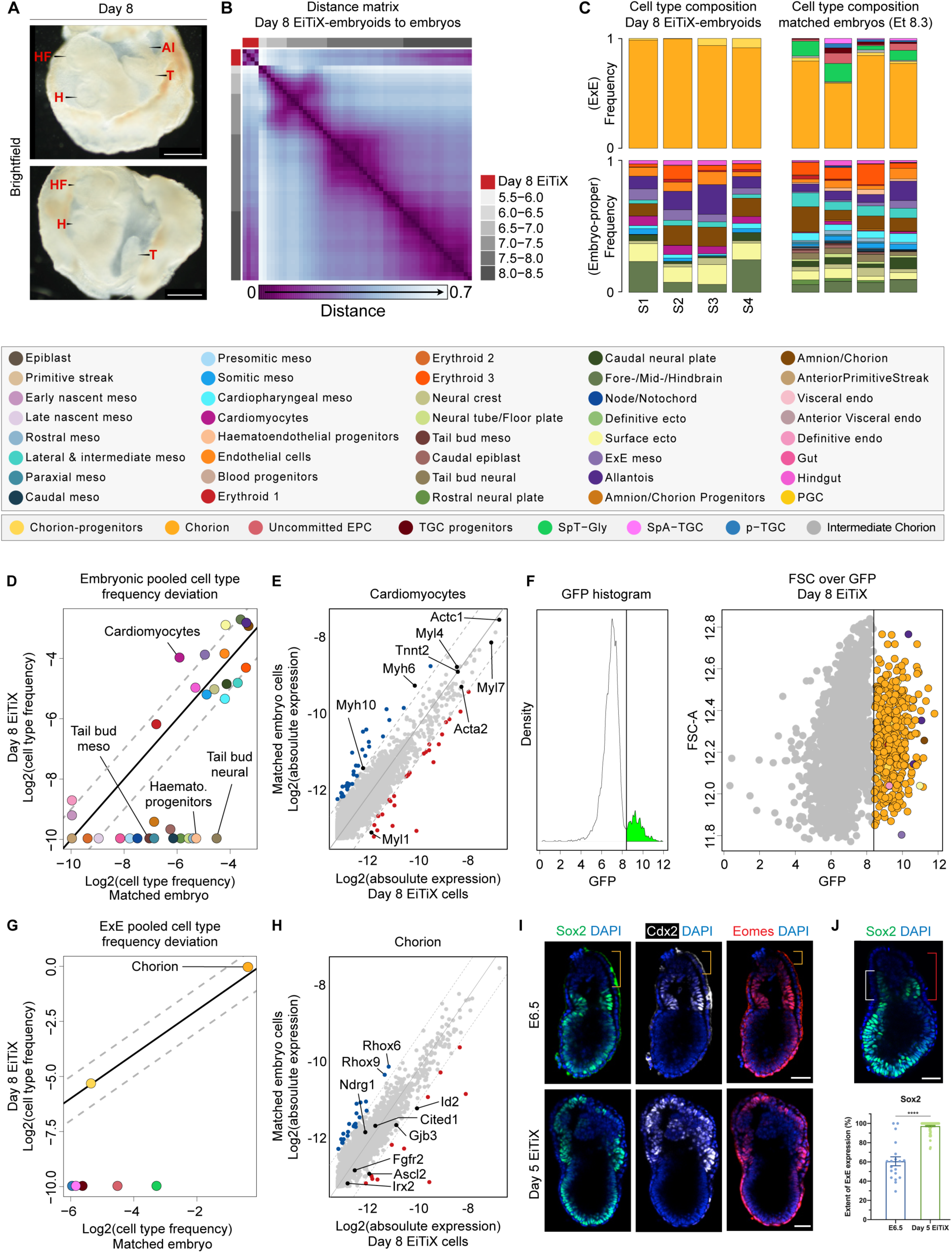
Cell state and composition analysis of neurulating embryoids using scRNA-seq. (**A**) Brightfield images of Day 8 EiTiX-embryoids collected for single-structure, single-cell RNA sequencing. The yolk sac-like membrane was partially opened to reveal embryonic structures. Al: allantois, H: heart, HF: headfolds, T: tailbud. Scale bar: 500µm. (**B**) Embryo-embryo cell type composition similarity matrix. Natural embryos are annotated based on embryonic age groups (grey), Day 8 EiTiX-embryoids (red). (**C**) Cell type composition bars of individual Day 8 EiTiX-embryoids (left) and matched natural embryos (right, annotated according to the legend below). (**D, G**) Pooled embryonic (D) and ExE (G) cell type frequencies comparison between Day 8 EiTiX-embryoids and matched natural embryos. (**E, H**) Bulk differential gene expression per cell state of Day 6 EiTiX cells against matched embryo cells in embryonic cell type (E, cardiomyocyte) and ExE cell type (H, chorion). Dots represent individual genes. Colour annotated dots mark genes with a two-fold change in expression (blue – above two-fold decrease in Day 6 EiTiX cells, red – above two-fold increase in Day 6 EiTiX cells). (**F**) GFP channel bimodal distribution (left) with threshold use to define GFP+ cell population shown as dots, annotated accordingly to cell type (right). (**I**) Day 5 EiTiX-embryoids and E6.5 natural embryos stained to reveal trophoblast stem markers Sox2 (green), Cdx2 (white) and Eomes (red). Orange brackets show the absence of trophoblast stem markers in the tip of ExE in E6.5 embryos. Scale bar: 50µm. (**J**) Quantification of the extent of ExE expression of Sox2. It was determined by dividing the height of the expression domain (white bracket) by the height of ExE (red bracket), multiplied by 100%. n = 19 E6.5 embryos from 2 experiments and 78 Day 5 EiTiX-embryoids from 3 experiments; ****p < 0.0001. Scale bar: 50µm.

Analyzing ExE differentiation showed expected progression in the chorion lineage (i.e. most chorion progenitors fully converting to their differentiated progenies). However, we could not detect any cell types associated with the EPC lineage, including uncommitted EPCs and trophoblast giant cells (TGC) progenitors (Figure 5C). We could confirm that the vast majority of GFP-positive i*Cdx2* ESCs (97.26%) gave rise to chorion lineage, although we noted a few embryonic cell types among the GFP positive cells (Figure 5F). Indeed, we occasionally observed GFP-positive cells in the EPI-like compartment in Day 4 EiTiX-embryoids, and we suspected that these cells might have retained ESC fate and would eventually give rise to embryonic lineages. Overall, cells in the chorion lineage appeared with comparable frequencies and exhibited highly similar gene expression signatures when compared to time-matched natural embryos (Figures 5G and 5H). Our inability to detect EPC and TGC subtypes in both Day 6 and Day 8 EiTiX-embryoids suggested i*Cdx2* ESCs exhibit restricted ExE differentiation potential. In the natural embryo, the ExE can be subdivided into proximal ExE (adjacent to the EPI/ExE boundary) and distal ExE (towards the tip of the embryo). The proximal ExE is characterized by the expression of TSC-like markers such as *Sox2*, *Cdx2* and *Eomes*, which are downregulated in the distal ExE, where the EPC and TGC subtypes are found (Donnison, Broadhurst and Pfeffer, 2015). We confirmed the absence of TSC-like markers in the distal ExE in E6.5 embryos (orange bracket), whereas in Day 5 EiTiX-embryoids, we observed strong expression of these genes throughout the ExE compartment (Figure 5I), with an extended-expression domain (Figure 5J and S5C).

## Discussion

Here we show that ESCs carrying a *Cdx2* transgene can adopt a TSC-like cell fate upon doxycycline-induced *Cdx2* overexpression. The resulting i*Cdx2* ESCs have the ability to self-assemble with ESCs induced to overexpress *Gata4* (induced XEN cells) and wildtype ESCs to generate an *in vitro* model of mouse post-implantation development with embryonic and extraembryonic lineages. The embryonic-extraembryonic embryo model we present here, which we term “EiTiX embryoids”, is derived entirely of ESCs and thus circumvents the use of undefined media to culture conventional extraembryonic cell lines. Such EiTiX-embryoids specify the DVE and AVE, establish an anterior-posterior axis, and undergo gastrulation. Following transfer into enriched *ex utero* culture media, EiTiX-embryoids undertake neurulation and form headfolds, brain, a beating heart structure, and develop extraembryonic tissues including yolk sac and chorion.

We performed single-structure scRNA-seq of 14 Day 6 and 4 Day 8 EiTiX-embryoids and projected the data on a temporal model describing the parallel differentiation in embryonic and extraembryonic lineages. This enabled direct comparisons with time-matched natural embryos, providing an analytical framework for quantifying the fidelity of intracellular transcriptional programs and overall cell composition within individual structures. We observed overall similarity of Day 6 EiTiX-embryoids with natural embryos of E6.0 to E7.5 stages, whereas Day 8 EiTiX-embryoids were most similar to natural embryos from E8.0 to E8.5. Despite the morphological variability of EiTiX-embryoids, transcriptional states appeared remarkably conserved compared to corresponding ones in natural embryos. We noted two main types of deviation from the natural flow of the embryo proper: (i) First, some differentiated cell types were missing in both Day 6 and Day 8 embryoids, and (ii) synchronicity between lineages was impaired in Day 6 embryos. Nevertheless, adaptations in culture conditions significantly improved lineage synchronicity in Day 8 embryoids, resulting in much comparable cell compositions to that of time-matched embryos. The analytical approach described here can complement future screening aimed at improved culture conditions, by providing a robust quantitative readout on embryoid development.

The EiTiX-embryo is thus able to develop many more tissues than structures derived solely from homogeneous populations of ESCs induced to differentiate by various exogenous molecules, generating a more complete *in vitro* model with both embryonic and extraembryonic tissues. Hence, our data substantiate the essential role of extraembryonic tissues in driving the self-organization of mouse embryo-like structures. For example, the role of ESCs induced to express Gata4 is of critical importance in establishing the formation of the AVE, which is required to direct the formation of anterior structures, particularly such as those of the forebrain.

Although the induced extraembryonic structures contribute to the correct development of diverse embryonic cell types and overall structure in EiTiX-embryoids, the development of the ExE lineage is incomplete as reflected by the lack of EPC and TGC cell types. This can be partly because EiTiX-embryoids lack the interactions with the maternal environment that they would have *in utero*; and partly because transcription factor-mediated induction biases *iCdx2* ESCs to differentiate into chorionic cell types. It is possible that there are two types of progenitor cells in the ExE splitting immediately after implantation and *iCdx2* ESCs resemble most the chorion lineage progneitors. Unlike the ExE in E6.5 natural embryos, we showed that there is a strong and extended expression of TSC-like markers throughout the ExE-like compartment in Day 5 EiTiX-embryoids. Thus, it might be necessary to introduce the expression of genes promoting ExE differentiation or induce the downregulation of TSC-like genes to generate EPC and TGC subtypes in EiTiX embryo. Moreover, by incorporating trophectoderm-derived cell types, our system offers future possibilities for dissecting the precise roles of such cells in the developmental process.

Despite not recapitulating the later stages of development of extraembryonic tissues, the substitution of TSCs by iCdx2 ESCs in EiTiX-embryoid permits remarkable development of the embryo *per se,* with the development of a yolk sac and chorion. The reconstitution of the three principal lineages of peri-implantation development exclusively from ESCs ensures simplified, defined and consistent culture conditions to recapitulating the interactions between embryonic and extraembryonic tissues and facilitate development through gastrulation to neurulation-like stages.

## Acknowledgements

The authors would like to thank David Glover and Ron Hadas for helpful comments, Yoav Mayshar for his assistance with sample collection for MARS-seq, Netta Reines for technical support with scRNA-seq processing, and the flow cytometry facility from the School of the Biological Sciences, University of Cambridge for their assistance in this work. The grants to M.Z.-G that supported this work are: NIH Pioneer Award (DP1 HD104575-01), European Research Council (669198), the Wellcome Trust (207415/Z/17/Z), Open Philanthropy/Silicon Valley Community Foundation and Weston Havens Foundation. The grants to YS that supported this work are: European Research Council (ERC_StG 852865) and Helen and Martin Kimmel Stem Cell Institute. K.Y.C.L. is supported by the Croucher Foundation and the Cambridge Trust. C.W.G. is supported by a Leverhulm Early Career Research Fellowship. Research in the M.Z.-G and Y.S. labs is supported by the Schwartz/Reisman Collaborative Science Program.

## Author contributions

Conceptualisation: G.A., Y.S., M.Z.-G.; Data curation: H.R.; Formal Analysis: K.Y.C.L., H.R.; Funding acquisition: Y.S., M.Z.-G.; Investigation: K.Y.C.L., C.W.G.; Methodology: K.Y.C.L., G.A.; Software: H.R.; Visualisation: K.Y.C.L., H.R.; Supervision: Y.S., M.Z.-G.; Writing: K.Y.C.L., H.R., Y.S., and M.Z.-G.

---

**Figure S1.**
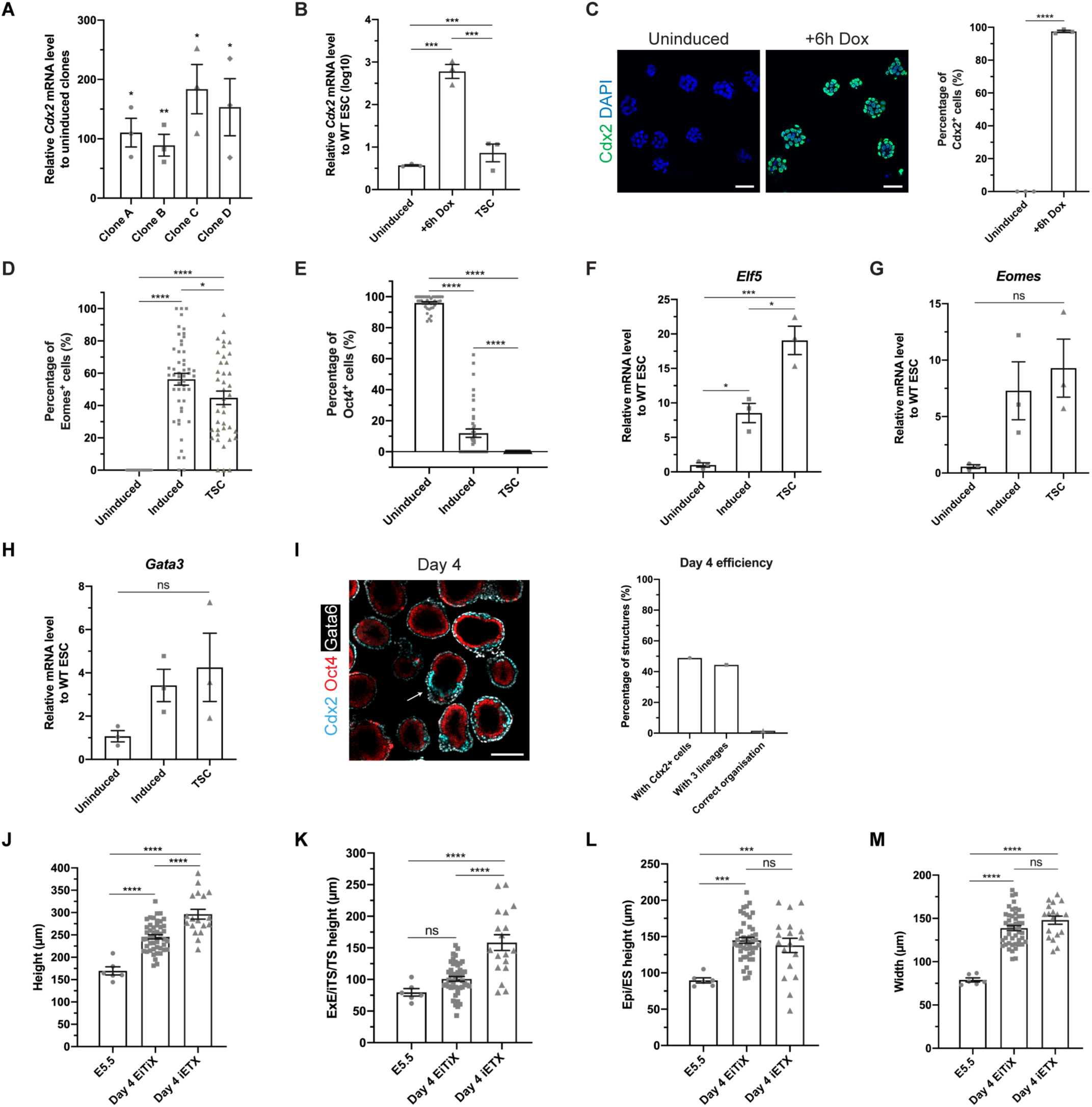
Characterisation of *Cdx2* inducible cells and Day 4 EiTiX-embryoids. (**A**) Fold changes of *Cdx2* mRNA level in four different clones of i*Cdx2* ESCs 6 hours after Dox induction, determined by qRT-PCR. Student’s t-test compares fold change with respect to uninduced counterparts. (**B**) Relative *Cdx2* mRNA levels in uninduced i*Cdx2* ESCs, 6-hour induced CAG iCdx2 ESCs, and TSCs as compared to unmodified ESCs. (**C**) Immunofluorescence images of Cdx2 expression in uninduced and 6-hour induced iCdx2 ESCs. Percentage of Cdx2^+^ cells is shown in righthand graph. n = 3 experiments, 5-10 random fields imaged for each condition in each experiment. Scale bar, 50µm. (**D-E**) Percentage of Eomes^+^ (D) and Oct4^+^ (E) cells in Day 3 aggregates of uninduced, induced i*Cdx2* ESCs and TSCs. (D) n = 43 uninduced aggregates, 50 induced aggregates and 39 TSC aggregates. (E) n = 40 uninduced aggregates, 40 induced aggregates and 43 TSC aggregates. (**F-H**) Relative mRNA levels of TSC markers *Elf5* (F), *Eomes* (G) and *Gata3* (H) in Day 3 aggregates, normalised to mRNA levels in unmodified ESC aggregates. (**I**) Day 4 structures generated using the iETX embryo protocol, stained to reveal Cdx2 (cyan), Oct4 (red) and Gata6 (white). Arrow indicates structure with correct organisation. Scale bar, 150µm. Graph shows percentages of structures with Cdx2-positive cells, with cells from all three lineages and with correct organisation. Scale bar, 150µm. (**J-M**) Size comparison of E5.5 mouse embryos, Day 4 EiTiX-embryoids and Day 4 iETX embryos in terms of height (J), ExE/iTS/TS height (K), Epi/ES height (L) and width (M). n = 6 E5.5 mouse embryos, 46 Day 4 EiTiX-embryoids and 15 Day 4 iETX embryos. All experiments were performed minimum 3 times with the exception of (I). Statistics: (A and C) Student’s t-test. (B, D-H, J-M) One-way ANOVA followed by Bonferroni’s multiple comparisons test. *p < 0.05; **p < 0.01; ***p < 0.001; ****p < 0.0001; ns, non-significant.

**Figure S2.**
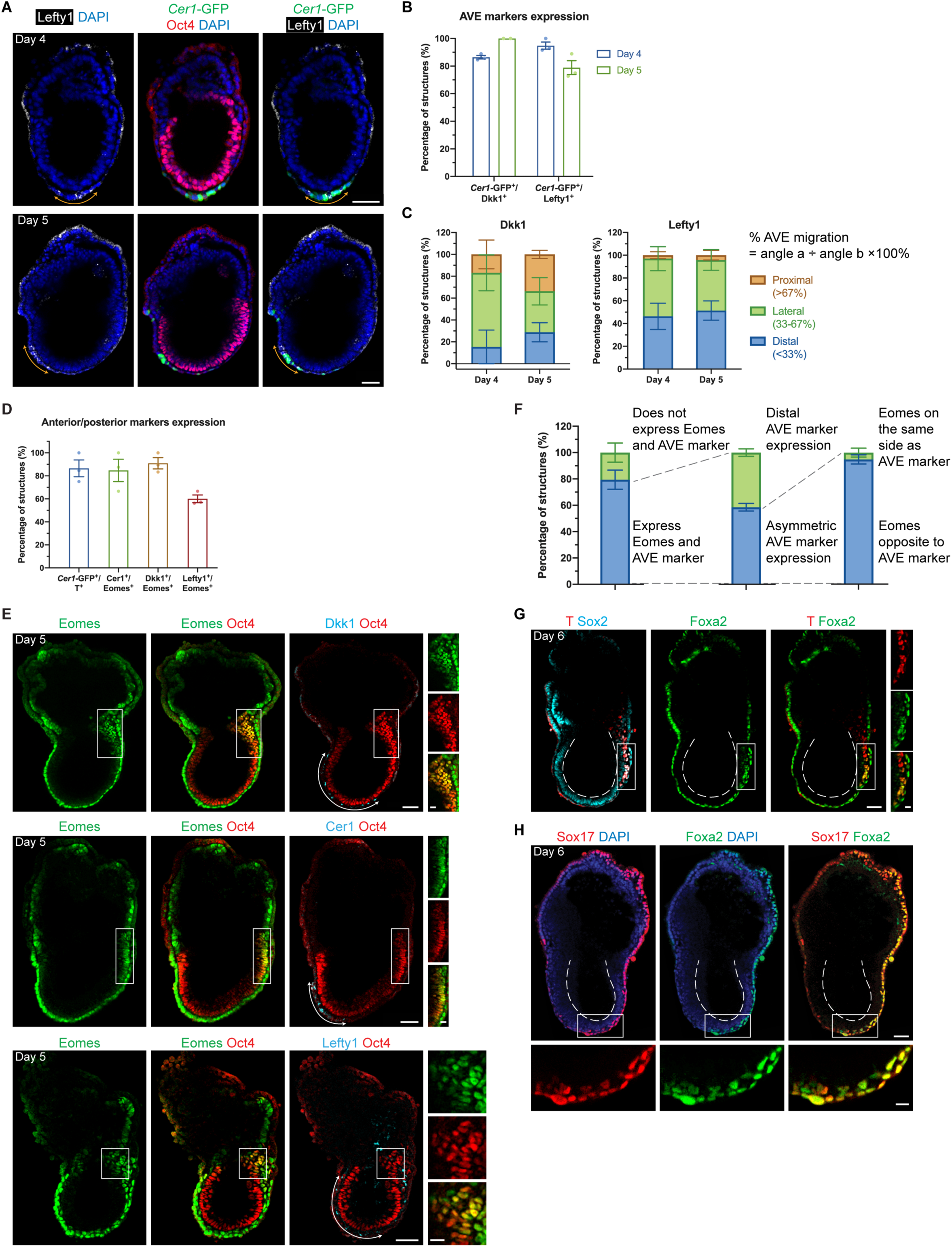
EiTiX-embryoids establish an anterior-posterior axis and undergo gastrulation. (**A**) Day 4 and Day 5 EiTiX-embryoids stained to reveal *Cer1*-GFP (green), Oct4 (red) and Lefty1 (white). Arrows indicate the Lefty1-positive domain. (**B**) Percentages of EiTiX-embryoids expressing different combinations of AVE markers. n = 29-32 Day 4 structures and 18-28 Day 5 structures. (**C**) Localisation of Dkk1 and Lefty1 in Day 4 and Day 5 EiTiX-embryoids. See Materials and Methods for quantification method. Dkk1: n = 29 Day 4 structures, 18 Day 5 structures; Lefty1: n = 30 Day 4 structures, 22 Day 5 structures from 3 experiments. (**D**) Percentages of EiTiX-embryoids expressing different combinations of anterior/posterior markers. n = 42 Day 5 structures (*Cer1*-GFP^+^/T^+^), 39 Day 5 structures (Cer1^+^/Eomes^+^), 37 Day 5 structures (Dkk1^+^/Eomes^+^) and 38 Day 5 structures (Lefty1^+^/Eomes^+^). (**E**) Day 5 EiTiX-embryoids stained for Eomes (green), Oct4 (red) and Cer1 or Dkk1 or Lefty1 (cyan). Arrows indicate Dkk1- or Cer1- or Lefty1-positive domain while the box indicates Dkk1- or Cer1- or Lefty1- and Eomes-double positive domains. (**F**) Graph shows percentages of Day 5 EiTiX-embryoids with 1) AVE marker and Eomes expression, 2) asymmetric AVE marker expression, and 3) Eomes expression on the opposite side to AVE. n = 114 structures. (**G**) Day 6 EiTiX-embryoid stained to reveal T (red) and Foxa2 (green). Box indicates T- and Foxa2- double positive domain; while dotted lines, lumen of ES compartment. n = 15/16 structures with T- and Foxa2- double positive cells. (**H**) Day 6 EiTiX-embryoid stained for Foxa2 (green) and Sox17 (red). Box, Foxa2- and Sox17- double positive domain; dotted lines, lumen of ES compartment. n = 15/17 structures with Foxa2- and Sox17-double positive cells. All experiments were performed minimum 2 times. Scale bars: 50µm, 15µm (zoomed).

**Figure S3.**
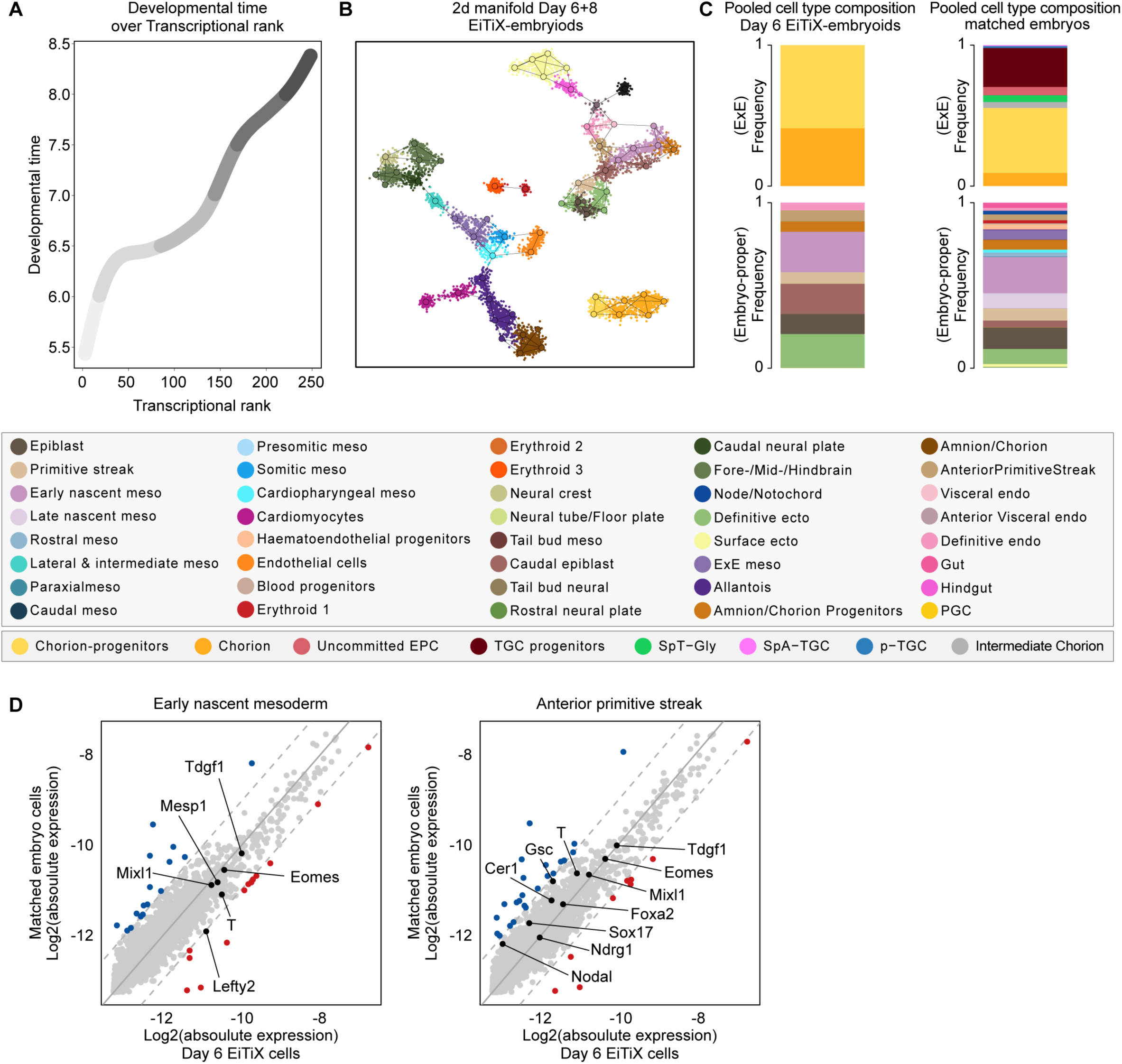
Day 6 EiTiX-embryoids capture major cell types of gastrulation. (**A**) Developmental time (E_t_) over embryo rank, annotated by age group (in 1E_t_ - 1.5E_t_ intervals, legend in Fig. 3C). (**B**) Day 6 and 8 EiTiX combined manifold, single cells (small dots) and Metacells (big dots) annotated by cell state (legend below). (**C**) Pooled ExE (top) and embryonic (bottom) cell-state frequencies of Day 6 EiTiX- embryoids (left panel) and time-matched natural embryos (right panel). (**C**) Bulk differential gene expression per cell state of Day 6 EiTiX cells against matched embryo cells; early nascent mesoderm (left) and anterior primitive streak (right). Dots represent individual genes. Colour annotated dots mark genes with a two-fold change in expression (blue – above two-fold decrease in Day 6 EiTiX cells, red – above two-fold increase in Day 6 EiTiX cells).

**Figure S4.**
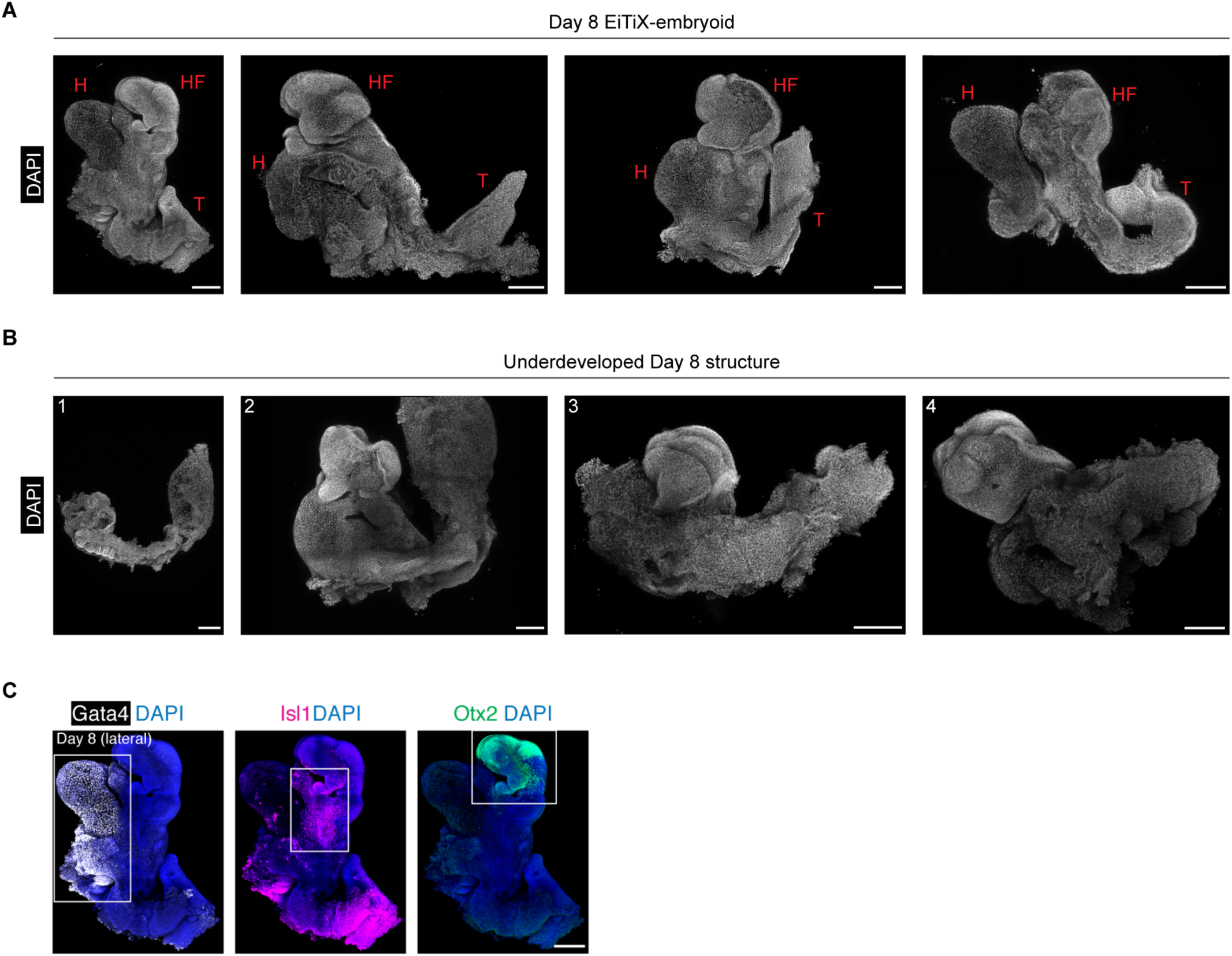
EiTiX-embryoids develop to late headfold stages with heart and chorion development. Examples of DAPI stained Day 8 EiTIX-embryoids (**A**) and underdeveloped Day 8 structures (**B**). Underdeveloped Day 8 structures showed stunted overall development (1) or impaired axial elongation to generate posterior structures (2-4). H: heart, HF: headfolds, T: tailbud. Scale bar: 200µm. (**C**) Lateral view of Day 8 EiTiX-embryoid stained to reveal heart marker Gata4 (white), pharyngeal mesoderm marker Isl1 (magenta), and forebrain marker Otx2 (green). Scale bar, 200µm

**Figure. S5.**
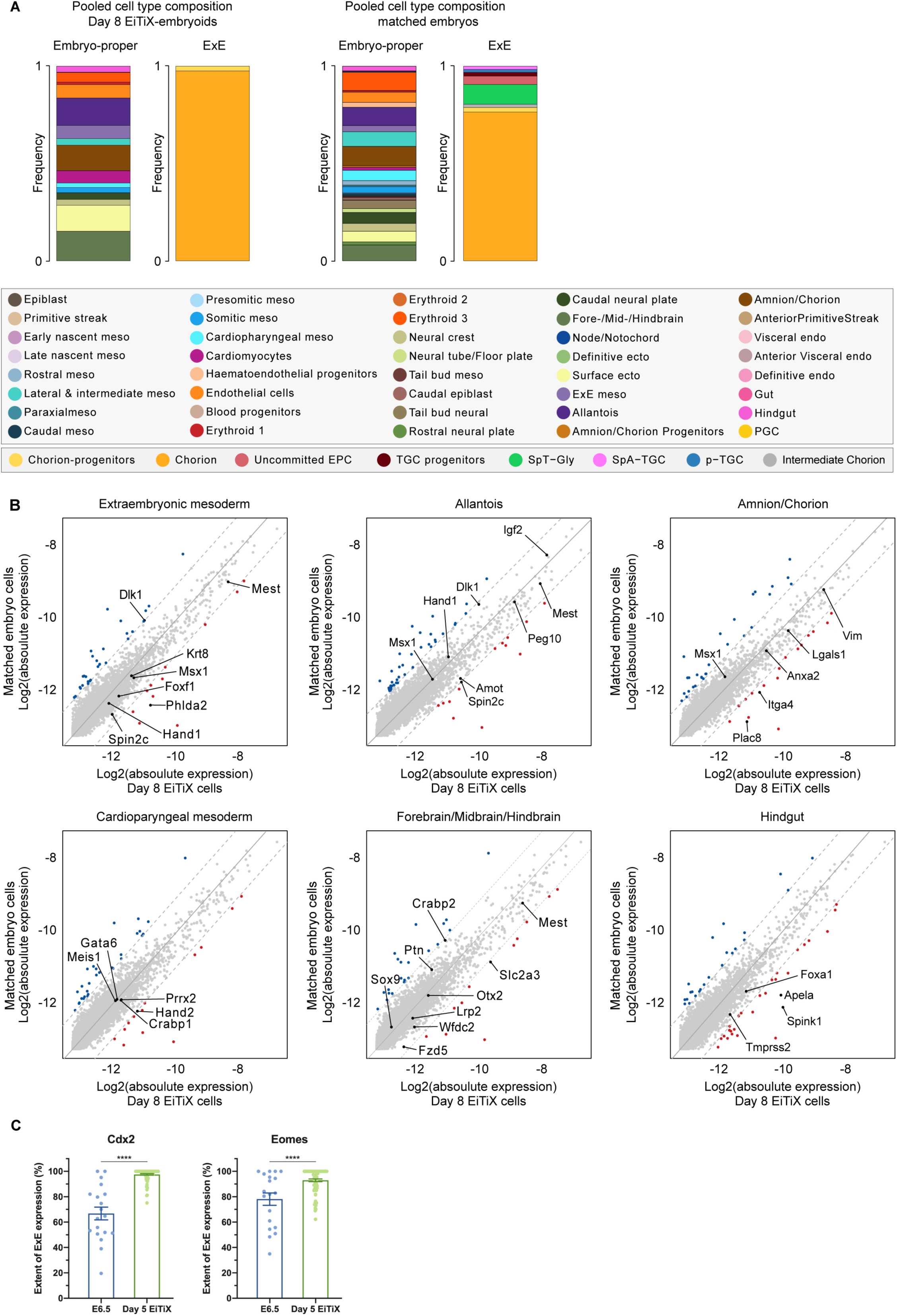
Cell state and composition analysis of neurulating embryoids using scRNA-seq. (**A**) Pooled ExE (left bar) and embryonic (right bar) cell-state frequencies of Day 8 EiTiX structures (left panel) and time-matched natural embryos (right panel, annotated according to the legend below). (**B**) Bulk differential gene expression per cell state of Day 8 EiTiX cells against matched embryo cells (extraembryonic mesoderm, allantois, amnion/chorion, cardiopharyngeal mesoderm, forebrain/midbrain/hindbrain and hindgut). Dots represent individual genes. Colour annotated dots mark genes with a two-fold change in expression (blue – above two-fold decrease in Day 8 EiTiX cells, red – above two-fold increase in Day 8 EiTiX cells). (**C**) Quantification of the extent of ExE expression Cdx2 and Eomes in E6.5 embryos and Day 5 EiTiX-embryoids. n = 19 E6.5 embryos from 2 experiments and 78 Day 5 EiTiX-embryoids from 3 experiments; ****p < 0.0001.

**Supplemental video**

**Video S1-2.** Video showing the beating heart region of Day 8 EiTiX-embryoids.

## Methods

### Cell culture

All cell lines were cultured at 37°C, 5% CO2 and 21% O2, and were passaged routinely every 3-4 days. Mycoplasma tests were carried out every two weeks to exclude contamination. ESCs were cultured and passaged as previously reported (Amadei *et al*., 2021).

### Cell lines

We have used the following ESC lines:

- CAG-GFP mouse ESCs: ESCs with constitutive membrane GFP expression, derived from CAG-GFP reporter mice (Rhee *et al*., 2006).
- CAG-GFP tetO-Cdx2 mouse ESCs: ESCs overexpressing Cdx2 upon Dox induction, generated in-house using methods described below.
- CAG-GFP tetO-Gata4 mouse ESCs: ESCs overexpressing Gata4 upon Dox induction generated as previously reported (Amadei *et al*., 2021)
- Cer1-GFP tetoO-Gata4 mouse ESCs: ESCs with GFP expression under Cer1 promoter which overexpress Gata4 upon Dox induction, generated as previously reported (Amadei *et al*., 2021)
- CD1 mouse ESCs: a generous gift from Jennifer Nichols
- CD1 tetO-Gata4 mouse ESCs: ESCs overexpressing Gata4 upon Dox induction generated as previously reported
- Confetti mouse TSCs: a generous gift from Jennifer Nichols
- mT/mG mouse ESCs: ESCs expressing membrane tdTomato, derived from mT/mG mice (Muzumdar *et al*., 2007)
- Sox2-Venus/Brachyury-mCherry/Oct4-Venus mouse ESCs: a generous gift from Jesse Veenvliet and Bernhard G. Hermann

### Plasmids and transfection

Cdx2 DNA flanked by attB sites was PCR-amplified from TSC cDNA using Cdx2-attB primers. Using Gateway technology (Thermo Fisher Scientific), it was then cloned into PB-tetO-hygromycin according to the manufacturer’s instructions and the resulting plasmid, PB-tetO-hygro-Cdx2 was verified by Sanger sequencing.

PB-tetO-hygro-Cdx2 was then transfected into 12,000 CAG-GFP ESCs together with pBAse and rtTA-zeocyin (0.25 μg each) using Lipofectamine 3000 Transfection Reagent (Invitrogen L3000001). Antibiotic selection was performed for 7 days with hygromycin (1:250; Gibco 10687010) and zeocyin (1:1000; InvivoGen ant-zn-1), followed by clonal expansion.

PB-tetO-hygro, pBAse and rtTA-zeocyin were generously gifted by Dr Jose Silva from the Stem Cell Institute (Cambridge, UK).

### RNA extraction and qRT-PCR

RNA was extracted from cell pellets using either Trizol Reagent (Invitrogen 15596-026) or RNeasy Mini Kit (Qiagen 74104). It was subsequently reverse transcribed into cDNA with M-MuLV reverse transcriptase (New England Biolabs M0253S). qRT-PCR was carried out using SYBR Green PCR Master Mix (Applied Biosystems 4368708) and StepOnePlus™ Real-Time PCR System (Applied Biosystems). ΔΔCt method was used to calculate fold change using GAPDH as endogenous control.

### Formation of cell aggregates and EiTiX-embryoids

Cell aggregates and EiTiX-embryoids were generated largely following the previously described method (Amadei *et al*., 2021) with the following modifications.

For cell aggregates, 38,400 6-hour induced iCdx2 ESCs, 19,200 uninduced iCdx2 ESCs, and 19,200 TSCs were plated into individual AggreWells. They were cultured in FC supplemented with 1 μg ml^-1^ heparin (Sigma-Aldrich H3149-25KU), and 25 ng ml^-1^ FGF4 (R&D Systems 7486-F4-025) for three days (with Y-27632 added for the first 24 hours only), changing media every day.

For EiTiX-embryoids, 38,400 6-hour induced iCdx2 ESCs, 6,000 6-hour induced iGata4 ESCs, 6,000 WT ESCs were pooled together and plated into each AggreWell. They were resuspended in in 1mL FC supplemented with 1 μg ml^-1^ heparin, 25 ng ml^-1^ FGF4 and 6.7nM Y27632 (STEMCELL Technologies 72304) and were added dropwise to each AggreWell. The culture condition for EiTiX-embryoids followed the previous protocol from Day 2.

### Inclusion criteria of EiTiX embryos

All EiTiX embryos were collected from AggreWell on Day 4 and their morphologies were examined under a dissection microscope. Structures were selected for analyses or further culture if 1) there were two distinct cellular compartments enclosed by a thin outer cell layer, and 2) there was a clear epithelialised ES compartment with a central lumen. For Day 5 and Day 6 structures, we selected structures that were elongated and had a thick epithelial cell layer in the ES compartment that resembled the EPI in natural mouse embryos.

### Culture of EiTiX-embryoids in *ex utero* culture media (EUCM)

EUCM was prepared as described in (Aguilera-Castrejon *et al*., 2021). It consists of 25% DMEM (Gibco 11880) with 1x Glutamax (Gibco 35050061), 100 units/ml penicillin/streptomycin (ThermoFisher 15140122) and 11 mM HEPES (Gibco 15630056), plus 50% rat serum (Charles River Laboratories) and 25% human cord serum (Cambridge Blood and Stem Cell Biobank). Rat serum and human cord serum were thawed at room temperature and heat-inactivated for 30 minutes at 56°C. After preparation, EUCM was filter-sterilised and equilibrated at 37°C for 1 hour. Each selected Day 5 EiTiX-embryoids were transferred to one well of 48-well multi-well plate for suspension culture (Greiner Bio-One 677102), with 250µl EUCM per well. On Day 6, 100µl EUCM was removed and 250µl fresh EUCM was added per well. On Day 7, EiTiX-embryoids were transferred to a rotating bottle culture chamber apparatus. Up to 3 EiTiX-embryoids were cultured in the same rotating bottle that contained 2ml EUCM supplemented with 3.0 mg/ml of D-Glucose (Sigma G8644).

### Mouse model and recovery of mouse embryos

CD-1 mice were maintained in the University of Cambridge’s University Biomedical Services Combined Animal Facility, adhering to national and international guidelines. Experiments were performed under the regulation of the Animals (Scientific Procedures) Act 1986 Amendment Regulations 2012 and were reviewed by the University of Cambridge Animal Welfare and Ethical Review Body (AWERB). Experiments were also approved by the Home Office.

Natural mating was performed with six-week-old CD-1 females and mouse embryos were recovered at embryonic days E6.5 by dissecting from the deciduae in M2 medium, as we described before (M. Zernicka-Goetz *et al*., 1997).

### Immunofluorescence

Samples were fixed with 4% paraformaldehyde for 20 minutes at room temperature and then washed three times with PBST (0.1% Tween-20 in PBS). Samples were permeabilised in 0.1 M glycine and 0.3% Triton X-100 in PBS for 30 minutes at room temperature. After washing with PBST for three times, primary antibodies diluted in blocking buffer (10% FBS and 0.1% Tween 20 in PBS) were added and the samples incubated overnight at 4°C. Primary antibodies were removed the following day, and samples were washed three times with PBST, before adding secondary antibodies and DAPI. After incubating overnight at 4°C, samples were washed three times with PBST and mounted in a glass-bottom dish for imaging.

### Dissociation of EiTiX-embryoids for MARS-seq

Individual Day 6 EiTiX-embryoids were dissected into four pieces with needles in PBS which were then dissociated with 70µl TrypLE Express Enzyme (Gibco 12604021) at 37°C for 15 minutes, pipetting up and down every 5 minutes. Dissociation was stopped by adding 500µl FC media supplemented with Y27632 (1:2,000) and DAPI (1:2,000). Day 8 samples were dissociated with 200µl TrypLE Express Enzyme and 800µl media was added to stop the dissociation. The cell suspension was subsequently filtered through a 40µm cell strainer (Merck CLS431750) and further diluted with 2ml FC media. It was then sorted by FACSAria III (BD Biosciences) using index sorting into 384-well plates.

### MARS-seq library preparation

Single-cell cDNA libraries were prepared as previously described (Cheng *et al*., 2021; Mittnenzweig *et al*., 2021) following the MARS-seq protocol. MARS-seq libraries were processed using NextSeq 500 or NovaSeq 6000. The output reads were processed following the MARS-seq2.0 protocol (Keren-Shaul *et al*., 2019) with the same specifications as previously reported, using STAR aligner for sequence alignment (Dobin *et al*., 2013). Here, we processed 8832 wells. To analyze Day 8 EiTiX embryoids, we used FACS index sorting to record fluorescence per cell in addition to structure identity per well. We then distinguished GFP positive cells using the green channel bimodal distribution.

### Atlas projection and cell type annotation of EiTiX-embryoids

To identify marker-genes for metacell construction (Baran *et al*., 2019), we selected all the genes displaying a minimal variance over mean (𝑇_𝑣𝑚 = 0.1) and coverage threshold (𝑇_𝑡𝑜𝑡_ = 50, 𝑇_𝑡𝑜𝑝_ = 3). These genes were clustered into 137 clusters based on their gene-gene correlation overall the UMI mat. Gene-cluster enriched with stress- and cell-cycle-related genes were manually removed (n = 625) leaving 807 feature genes. The final metacell object (𝐾𝑛𝑛 = 100, minimal metacell size = 30 cells) contained 60 metacells comprised of 7076 cells (2184 from Day 6 EiTiX structures and 4892 from Day 8 EiTiX-embryoids) with a 5362 median UMIs per cell. Metacells were annotated with cell types by projection on the gastrulation wildtype atlas, as previously reported (Cheng *et al*., 2021; Mittnenzweig *et al*., 2021).

### Natural embryo matching

For each EiTiX-embryoid, we inferred a best-matching natural embryo based on the similarity (Euclidean distance) between their cell state compositions. Only natural embryos with at least 161 embryonic cells and 29 extraembryonic ectoderm cells were included (threshold fits the calculated median number of ExE cells per EiTiX-embryoid). For each EiTiX-embryoid, we included the three closest natural embryos in the matching natural cohort, counting each natural embryo only once. Natural embryos were temporally ordered as previously reported (Cheng *et al*., 2021; Mittnenzweig *et al*., 2021).

### Mean differential expression among transcriptional states from EiTiX-embryoids

For each EiTiX Day, we computed bulk (average) gene expression profiles per cell type and compared them with the corresponding natural embryo expression profiles from the matching natural embryos (log2 absolute expression). For each cell type, the number of included natural embryo cells was down sampled to the corresponding number of cells from EiTiX-embryoids compared. Cell-cycle and stress-related genes were not included in that comparison. Highlighted cell-type-specific genes for each included cell type were defined as being on average at least two-fold enriched in the metacells from this cell type relative to the global average among all natural embryo Metacells and additional known cell type markers were added.

### Image acquisition, processing and analysis

Leica SP5 and SP8 confocal microscopes (Leica Microsystems) with either a 40x oil objective or a 25x water objective were used to acquire immunofluorescence images. A 405 nm diode laser, 488 nm argon laser, 543 nm HeNe laser and 633 nm HeNe laser (Alexa Fluor 647) were used to excite the fluorophores. Fiji and Smart Denoise (Gurdon Institute) were used for image processing and analysis.

### Quantification of the extent of AVE anterior localization

The angle between the distal tip and the most anterior cell expressing AVE marker was termed as angle a (white) while the angle between the distal tip and the boundary of Oct4-positive domain was termed as angle b (orange) (Figure 2D). % AVE migration was obtained by dividing angle a by angle b and multiplying by 100%. It was then classified as proximal (>67%), lateral (33-67%) and distal (<33%).

### Quantification of the extent of Brachyury (T) extension

The angle between the posterior boundary of Oct4-positive domain and the most anterior T-positive cell was termed as angle a (white) while the angle between the posterior boundary of Oct4-positive domain and the distal tip The percentage of T extension was obtained by dividing angle a by angle b and multiplying by 100%.

### Statistics

Statistical analyses were performed using GraphPad Prism 8 and quantitative data were presented as mean ± S.E.M. or as violin plots with median and quartiles. Student’s t-test was used to determine statistical significance between two samples while one-way ANOVA followed by Bonferroni’s multiple comparisons test was used to determine statistical significance between more than two groups. Sample size and number of experimental replicates were indicated in figure legends.

## Materials & Correspondence

All data are available upon request to the corresponding author. Codes used in the study and raw data of scRNA-seq can be found in https://github.com/hernanRubinstein/EiTiX-embryoids

